# Characterization of the SARS-CoV-2 BA.5 Variants in H11-K18-hACE2 Hamsters

**DOI:** 10.1101/2024.02.19.581112

**Authors:** Mei Dong, Haofeng Lin, Margaret Pan, Minghong Huang, Meiqin Liu, Rendi Jiang, Yana Lai, Aimin Shi, Bing Yao, Ben Hu, Zhengli Shi, Aihua Zhang, Yun Gao, Wentao Zeng, Li Jianmin

**Author notes:** Paste corresponding author name(s) here. **Email:** Corresponding author;). **Author Contributions:** Mei Dong, Haofeng Lin, Margaret Pan, Minghong Huang, Meiqin Liu contributed equally. **Competing Interest Statement:** Authors declare that they have no competing interests.

## Abstract

This study aims to comprehensively characterize the SARS-CoV-2 BA.5 variants using K18 hACE2 transgenic mice and golden hamsters as model organisms. Previous research on SARS-CoV-2 has utilized both mouse and hamster models, leading to conflicting results concerning the virus’s lethality. In our study, the finding suggests that H11-K18 hACE2 golden hamsters closely mimic the disease progression observed in human COVID-19 cases caused by BA.5 variants, demonstrating consistent severity and symptoms comparable to severe infections.

Additionally, hamsters exhibit heightened respiratory viral replication, accurately reflecting the clinical viral kinetics observed in humans. The study emphasizes the critical importance of selecting an appropriate animal model for SARS-CoV-2 research, while also providing robust support for the hypothesis that BA.5 variants contribute to fatal outcomes in COVID-19 cases. These findings highlight the pivotal role of the golden hamster model in advancing our understanding of the pathogenic mechanisms underlying SARS-CoV-2 variants, as well as in the development of targeted therapeutic strategies.

**Significance Statement:** Our research work explores groundbreaking insights that could reshape our understanding of COVID-19 and pave the way for targeted therapies. We use golden hamster models to express the possibility of different animal models could contribute to human diseases. We hope this finding could clarify some conflicts existed, and help further development of medication for COVID.

## Introduction

Severe Acute Respiratory Syndrome Coronavirus 2 (SARS-CoV-2) is the causative agent of coronavirus disease 2019 (COVID-19), a febrile respiratory illness in humans that emerged in late 2019 in China and subsequently spread worldwide[1,2].

Notably, SARS-CoV-2 infections can potentially result in multi-organ injuries and chronic diseases beyond the infectious stage. Various factors, including antiviral treatment and immune stress, lead to mutations across the entire genome of SARS-CoV-2, resulting in amino acid substitutions within its proteins. Several SARS-CoV-2 variants have emerged due to these amino acid substitutions.

The Omicron lineage of SARS-CoV-2 was first identified in November 2021, rapidly spreading and becoming globally dominant, subsequently dividing into several sublineages[3]. In early April 2022, a new Omicron lineage, designated as BA.5, was reported in Gauteng, South Africa. The BA.5 receptor-binding domain (RBD) exhibits enhanced ACE2 binding and evasion of immune recognition, suggesting that BA.5 may possess higher transmissibility compared to other Omicron variants[4,5]. In the currently known human-infecting SARS-related coronaviruses, serotypes of Omicron subvariants have been a new serotype independent of SARS-CoV-1, and its ancestral SARS-CoV-2. The diminishing efficacy of COVID-19 therapeutics underscores the significance of utilizing animal models to assess the potential of new treatments in mitigating the replication and transmission of both existing and emergent SARS-CoV-2 variants.

Mouse ACE2 has a low affinity for the prototype SARS-CoV-2 spike protein; hence, mice are naturally less susceptible to infection by ACE2-utilizing human coronaviruses[6–8]. In contrast to mice, hamsters are naturally susceptible to SARS-CoV-2 infection[9,10]. Previous investigations have shown a degree of controversy as attempts to replicate BA.5 variant dynamics in murine models yielded disparate findings. Consequently, two conjectures have emerged: one positing the potential lethality of BA.5 and the other suggesting its non-lethal attributes[11–13].

Transgenic mice are typically generated using either random or site-specific integration techniques. However, the random insertion of genes can lead to a variable phenotype, gene silencing, and unexpected expression due to position effects. The H11 locus is situated in an intergenic region near chromosome 11 centromere, flanked by Eif4enif1 and Drg1 genes in mice. As the H11 locus lacks an endogenous promoter, the integrated genes are not anticipated to disrupt any endogenous genes. The mouse models exhibit normal fertility and phenotype.

Through a comparative genomic analysis involving mice and golden hamsters, our investigation revealed the presence of the H11 locus within the golden hamster’s genome, positioned between the Eif4enif1 and Drg1 genes at the chromosome 4 centromere (Figure 1a)[14]. This structural similarity suggests that the H11 locus construct of the golden hamster resembles that of the mouse. Hence, we hypothesized that this similarity makes the H11 locus suitable for the expression of transgenes in the golden hamster. Specifically, we evaluated the efficacy of the mouse model featuring ectopic expression of the human Angiotensin-Converting Enzyme 2 (hACE2) receptor under the regulation of the Keratin 18 (K18) promoter, examining both fixed and random integration scenarios. Similarly, we assessed the hamster model with ectopic expression of the hACE2 receptor. The outcomes of this analysis provide valuable insights into the biological behavior of these variants, shedding light on their potential implications in the progression of severe COVID-19. Given the extensive morbidity and mortality, developing vaccines, antibody-based countermeasures, and antiviral agents has become a global health priority.

**Figure 1.**
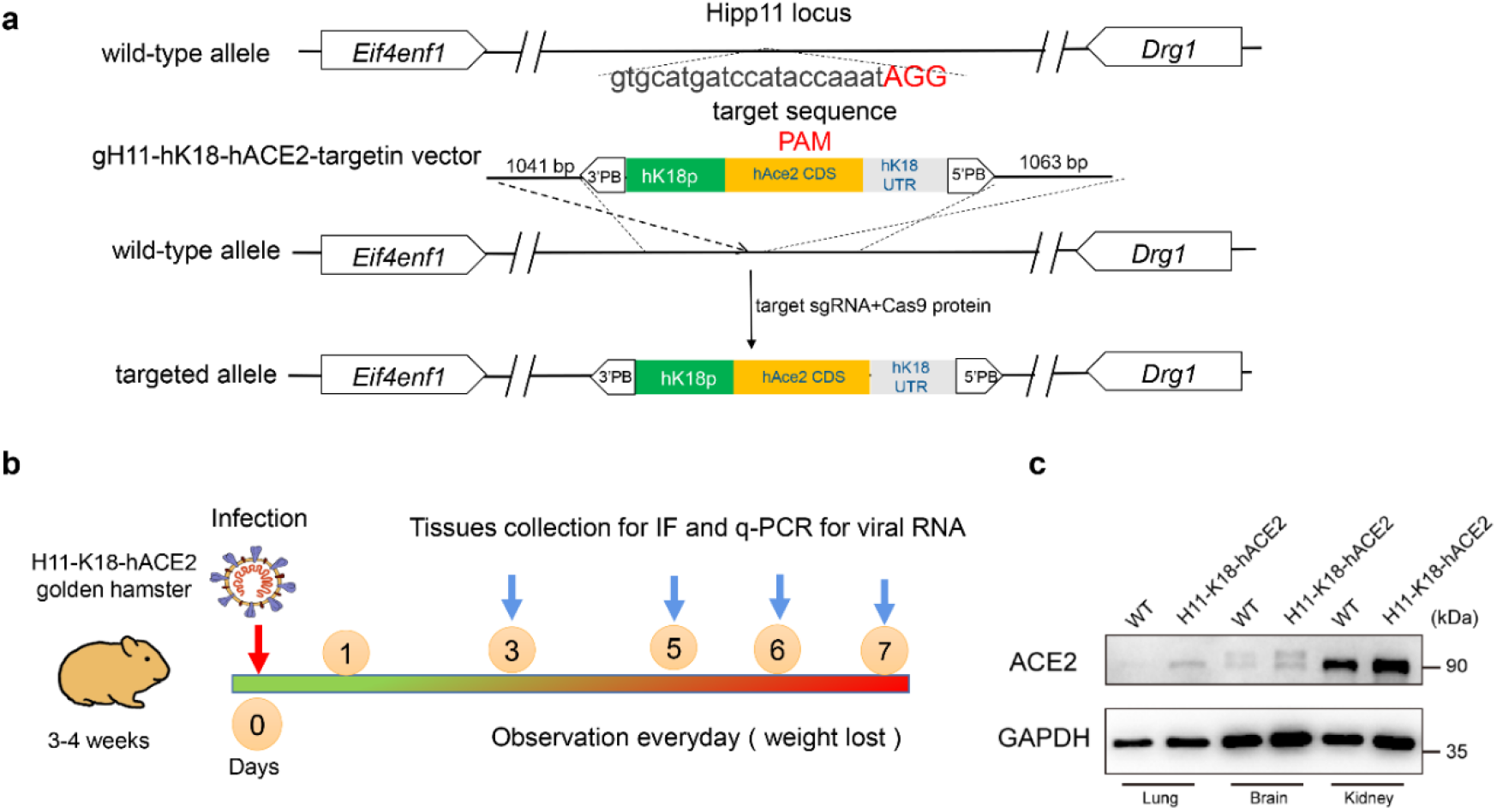
Experiment Scheme in SARS-CoV-2-Infected H11-K18-hACE2 golden hamster. **(a)** Schematic representation of H11-K18-hACE2 transgenic golden hamster by the Cas9-CRISPR technology. **(b)** H11-K18-hACE2 golden hamster were intranasally infected with 3 × 104 TCID50 COV1D-19 BA.5 virus each and sacrificed to collect tissue samples at 3, 5, 6 and 7 days post-infection (DPI). Golden hamster body weights were monitored for up to 7 days. Infected golden hamster that died showed noticeable body weight decreases from day 5 to 6 days. **(c)** Western blot analysis of ACE2 protein expression in kidney and lung tissues of wild-type (WT) and H11-K18-hACE2 transgenic golden hamsters.

## Results

### Generation of transgenic H11-K18-hACE2 golden hamster

The H11 locus in the golden hamster is situated in an intergenic region between Eif4enif1 and Drg1 (Figure 1a)[15]. The H11-K18-hACE2 transgenic golden hamster strains were generated by injecting a two-cell embryo with CRISPR–Cas9 using H11 sgRNA and the H11-K18-hACE2 donor plasmid[16,17]. The donor plasmid contained the K18-hACE2 cassette flanked by two 1kb H11 homologous arms (Figure 1b).

Microinjection followed by embryo transfer resulted in a cohort of 22 pups. HR-mediated K18-hACE2 cassette insertion was confirmed positive in two pups through PCR results. The humanized golden hamster model demonstrated increased expression levels of ACE2 protein compared to the wild-type (WT) in lung, brain, and kidney, as observed through Western blot analysis. This finding suggests that the humanized golden hamster model may be more suitable for studying various diseases and conditions related to these organs, such as viral infections, respiratory diseases, and neurological disorders. The enhanced expression levels in these tissues could potentially contribute to a better understanding of the underlying mechanisms and the development of targeted therapeutic strategies (Figure 1c). These data provide evidence for our site-specific integrase-mediated transgenesis in golden hamsters. Notably, the resulting offspring with both H11-K18-hACE2 heterozygotes and homozygotes followed the expected Mendelian inheritance ratio, exhibiting robust physiological health and reproductive capability, a trend consistently observed across the generated pups.

### H11-K18 hACE2 golden hamsters infected by BA.5 variants were lethality

Infection with BA.5 in mice and hamsters was characterized to assess the BA.5 variants in vivo. The BA.5 variants were isolated in VeroE6/TMPRSS2 cells through our investigation. Initially, we evaluated the pathogenicity of hACE2 mice with two different insertion modes against BA.5 isolates. Notably, significant body weight loss was observed from the first day post infection (dpi) following intranasal exposure to BA.5 variants in H11-K18-hACE2 mice and Tg-K18-hACE2 mice (Fig. 2a). In H11-K18-hACE2 mice, BA.5 infection led to 100% mortality by 5 dpi, while Tg-K18-hACE2 mice exhibited 100% mortality by 6 dpi (Fig. 2b). These findings are inconsistent with prior reports[18]. While hamster ACE2 can function as a receptor for the SARS-CoV-2 spike protein, some residues in contact with hACE2 are not well-preserved, potentially resulting in incomplete detection of infectivity and impacting result accuracy. To better simulate human infections, we developed transgenic hamsters named H11-K18-hACE2 hamsters, expressing hACE2 under the epithelial cytokeratin-18 promoter. Significant weight loss followed intranasal inoculation with the BA.5 variants (Fig. 1a), with half of the hamsters succumbing at 4 dpi and the rest at 5 dpi (Fig. 2b).

**Figure 2.**
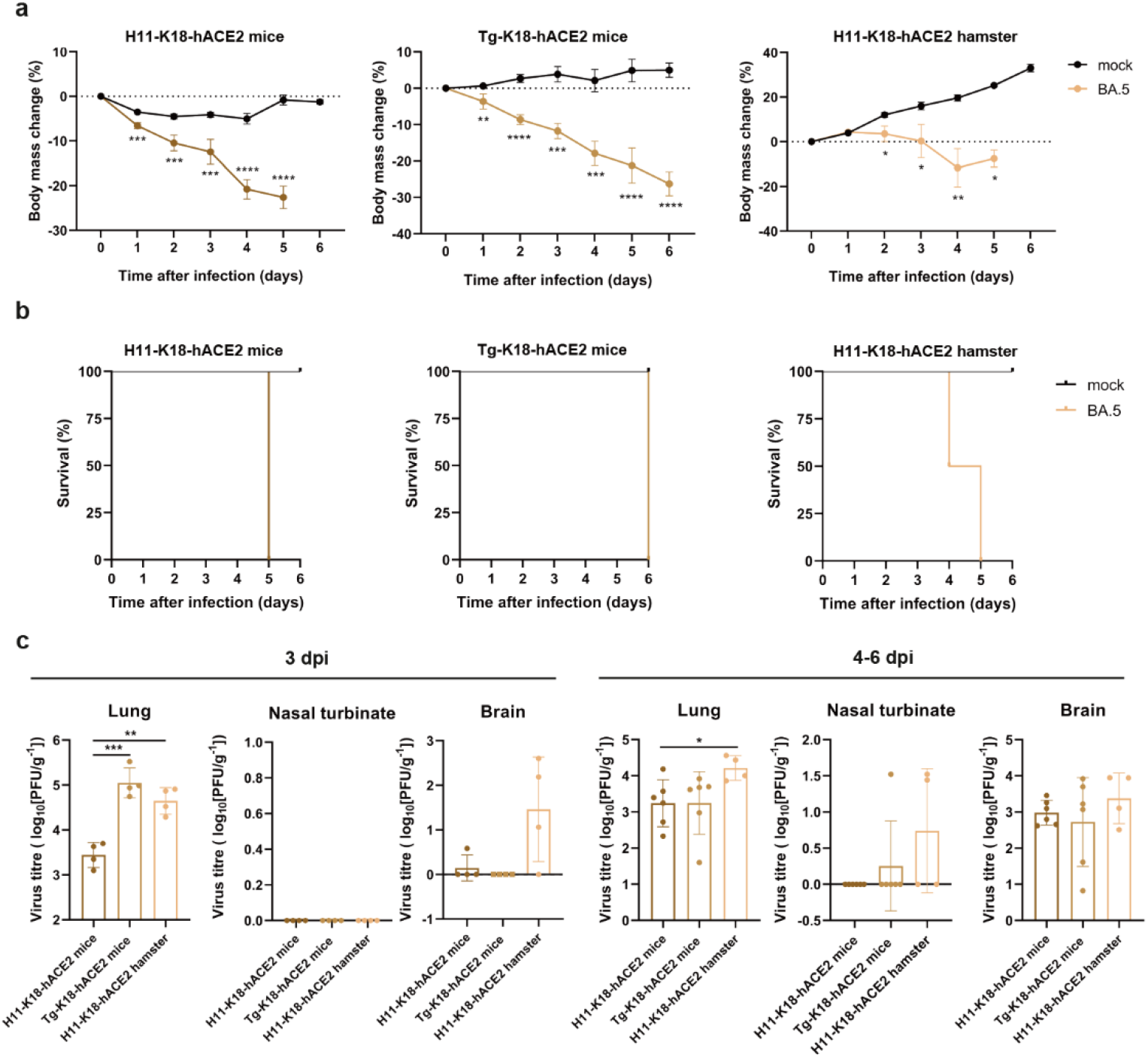
The replication and pathogenicity of BA.5 isolates in K18 hACE2 mice and hamsters. **(a-c)** H11-K18-hACE2 mice, Tg-K18-hACE2 mice and H11-K18-hACE2 hamsters were inoculated intranasally with 104 PFU in 50 μl BA.5 or PBS (mock). Body mass change **(a)** and survival **(b)** were monitored daily for 6 days. (mock, n=3; H11-K18-hACE2 mice, n=6; Tg-K18-hACE2 mice, n=6; H11-K18-hACE2 hamsters, n=4) **(c)** Virus replication in infected H11-K18-hACE2 mice, Tg-K18-hACE2 mice and H11-K18-hACE2 hamsters. Infected mice and hamsters were euthanized at 3 dpi and natural death for virus titration. Data represent means ± SEM (3 dpi, n=4; natural death, n=6). *p< 0.05, **p< 0.01, *** p< 0.001. Points indicate data from individual rodents. Difference between treatments was tested by a two-sample student t-test.

We assessed infection levels in the respiratory tract and brain of K18-hACE2 mice and hamsters (Fig. 2c). At 3 dpi, BA.5 virus replicated in the lungs of all infected animals. Virus titers were significantly higher in Tg-K18-hACE2 mice and H11-K18-hACE2 hamsters compared to H11-K18-hACE2 mice. However, no virus titers were detected in the nasal turbinates of either mice or hamsters infected with BA.5 at 3 dpi. Both BA.5-infected H11-K18-hACE2 mice and H11-K18-hACE2 hamsters exhibited comparable viral titers in the brain. Conversely, Tg-K18-hACE2 mice showed no detectable viral titers in the brain. After the natural deaths of these models, autopsies were conducted to determine virus titers. Both H11-K18-hACE2 mice (5 dpi) and Tg-K18-hACE2 mice (6 dpi) exhibited similar titers in their lungs, although H11-K18-hACE2 hamsters (4-5 dpi) had markedly elevated titers. Moreover, minor differences in infection were observed in the nasal turbinates of Tg-K18-hACE2 mice and H11-K18-hACE2 hamsters at the time of natural death. Conversely, H11-K18-hACE2 mice showed no detectable infection in the nasal turbinates. In summary, all BA.5-infected mice and hamsters exhibited a relatively preserved upper respiratory tract but impaired lower respiratory tract function. However, virus titers were higher at the time of death compared to 3 dpi in all mice and hamsters, demonstrating similar replicative abilities of BA.5 in the brains of both animal models.

### Histopathology in Infected Mice and Hamsters

To further understand the pathogenesis of SARS-CoV-2 infection in these animal models, histopathological analysis of the lungs and brains of H11-K18-hACE2 mice, Tg-K18-hACE2 mice, and H11-K18-hACE2 hamsters infected with BA.5 was conducted.

Comparable samples from healthy rodents were utilized as negative controls. At 3 dpi, both H11-K18-hACE2 mice and Tg-K18-hACE2 mice showed minimal inflammatory cell infiltration around the bronchi and bronchioles. In H11-K18-hACE2 hamsters, some minor changes were observed in lung tissues, including multifocal lesions and thickening of certain alveolar walls with minor amounts of fibrin exudation(Figure 3). Small foci of inflammatory cell infiltration into the alveolar spaces were noticed in H11-K18-hACE2 mice and Tg-K18-hACE2 mice at 5-6 dpi. H11-K18-hACE2 hamsters, on the other hand, exhibited fibrous tissue proliferation, obstruction of some terminal fine bronchi, and lysis and necrosis of some alveolar cells leading to mortality. However, there were no significant pathological differences observed in the brains of all mice and hamsters (Figure 4).

**Fig. 3.**
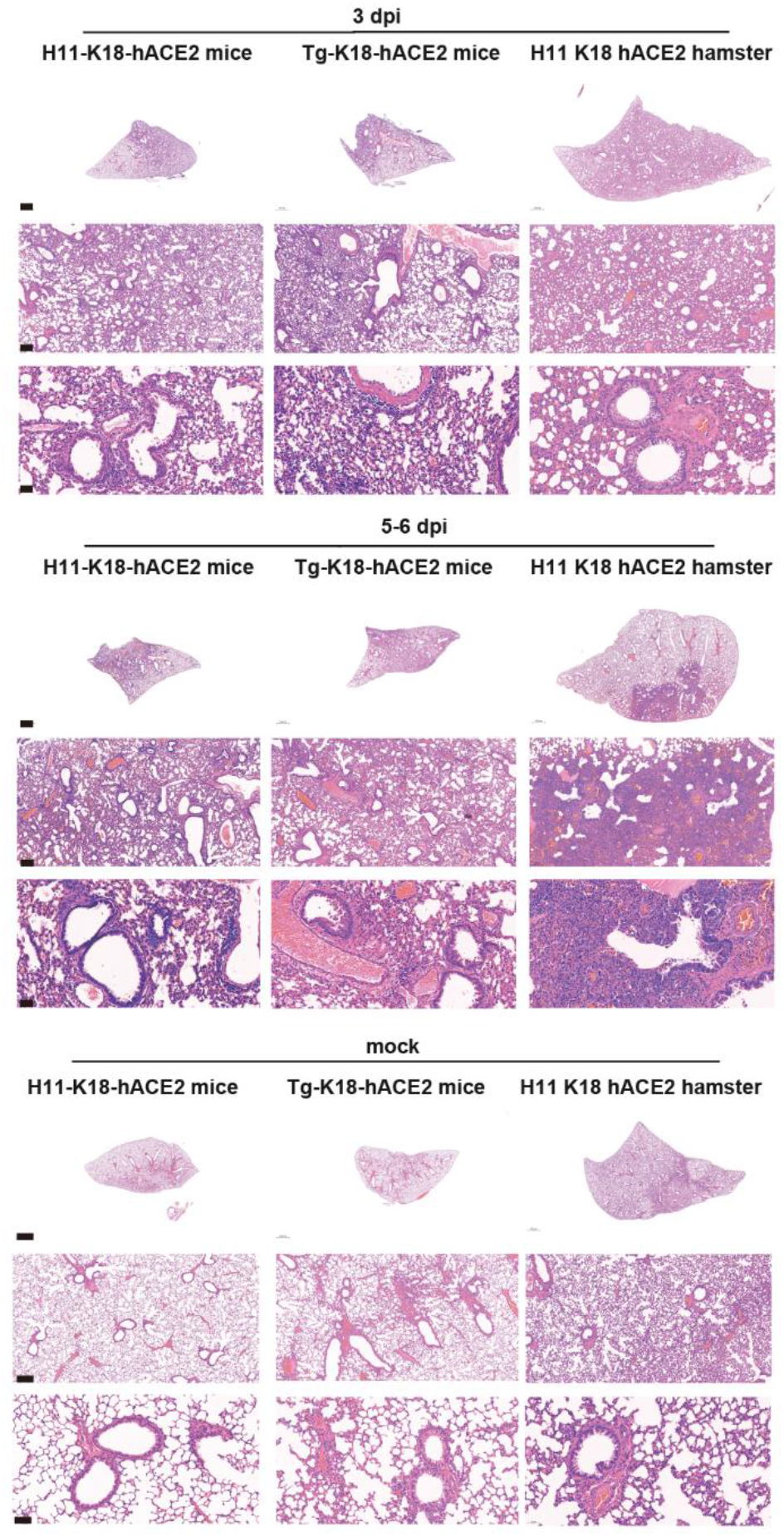
Histopathological phenotypes in the lungs of mice and hamsters inoculated with SARS-CoV-2 BA.5 variant. Euthanized mice and hamsters (3-4 each) were used to examine the pathological changes in the lungs 3 dpi, respectively. Natural death mice and hamsters (6 each) were used to examine the pathological changes in the lungs 5-6 dpi.

**Fig. 4.**
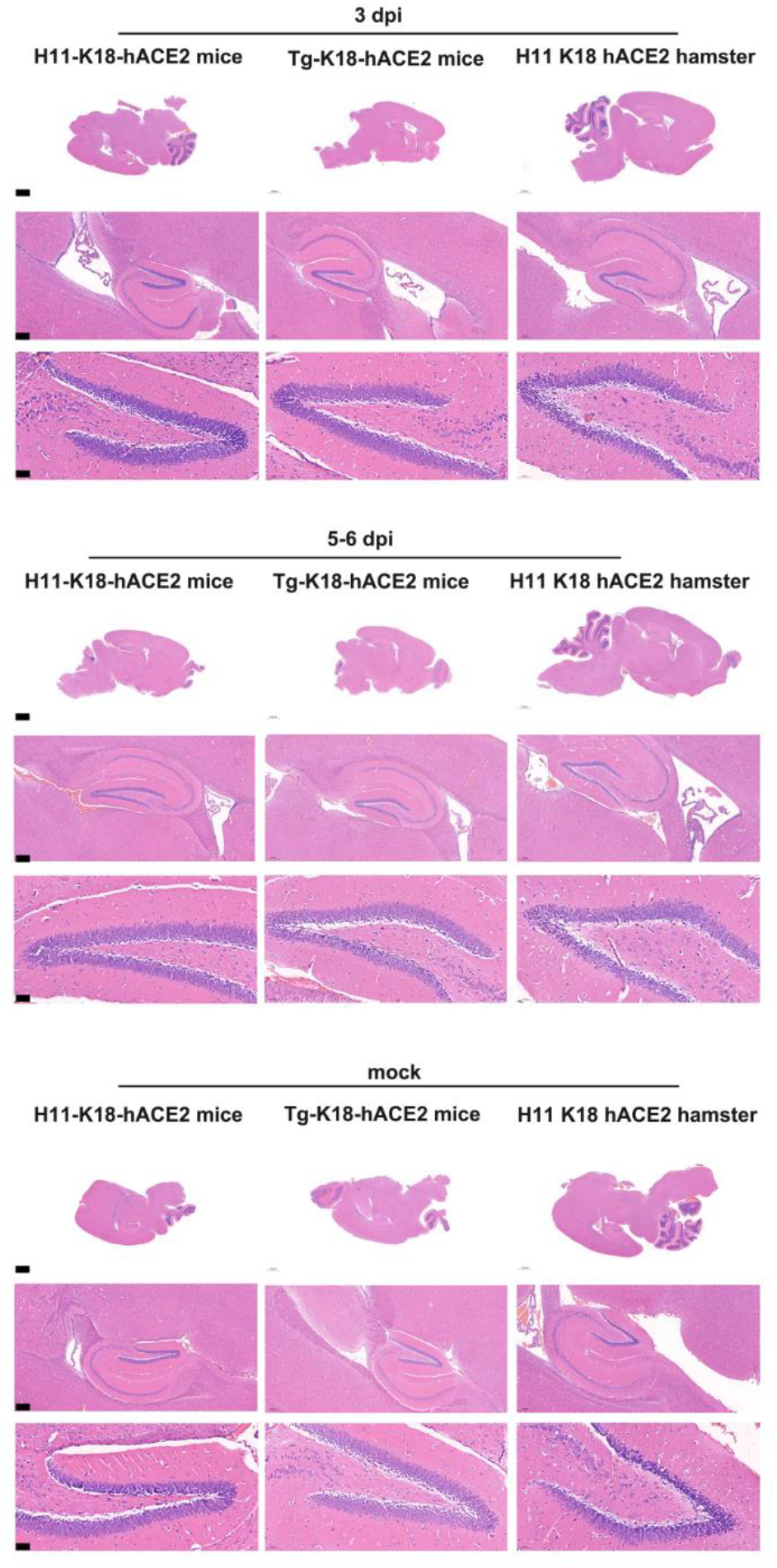
Histopathological phenotypes in the brains of mice and hamsters inoculated with SARS-CoV-2 BA.5 variant. Euthanized mice and hamsters (3-4 each) were used to examine the pathological changes in the brains 3 dpi, respectively. Natural death mice and hamsters (6 each) were used to examine the pathological changes in the brains 5-6 dpi.

### Immunofluorescence Analysis in Infected Mice and Hamsters

BA.5 was analyzed as low pathogenicity but enhanced virus spread/ inflammation compared with the ancestral strain[19]. To investigate the in vivo pathogenicity of BA.5 isolates and to elucidate the expression patterns of the SARS-CoV-2 viral nucleoprotein (NP) protein and ACE2 receptor in infected mice and hamsters, immunofluorescence analysis was conducted. The findings revealed distinct expression of NP proteins predominantly within the bronchioles of H11-K18-hACE2 hamsters (Figure 5). In contrast, BA.5-inoculated mice and hamsters, upon natural death, displayed SARS-CoV-2 NP protein presence in the bronchial and bronchiolar epithelium. Notably, accumulating evidence underscores SARS-CoV-2’s capacity to impact both the respiratory and central nervous systems[20,21]. In line with previous studies, our results depicted the presence of SARS-CoV-2 viral NP protein in the brain (Figure 6).

**Fig. 5.**
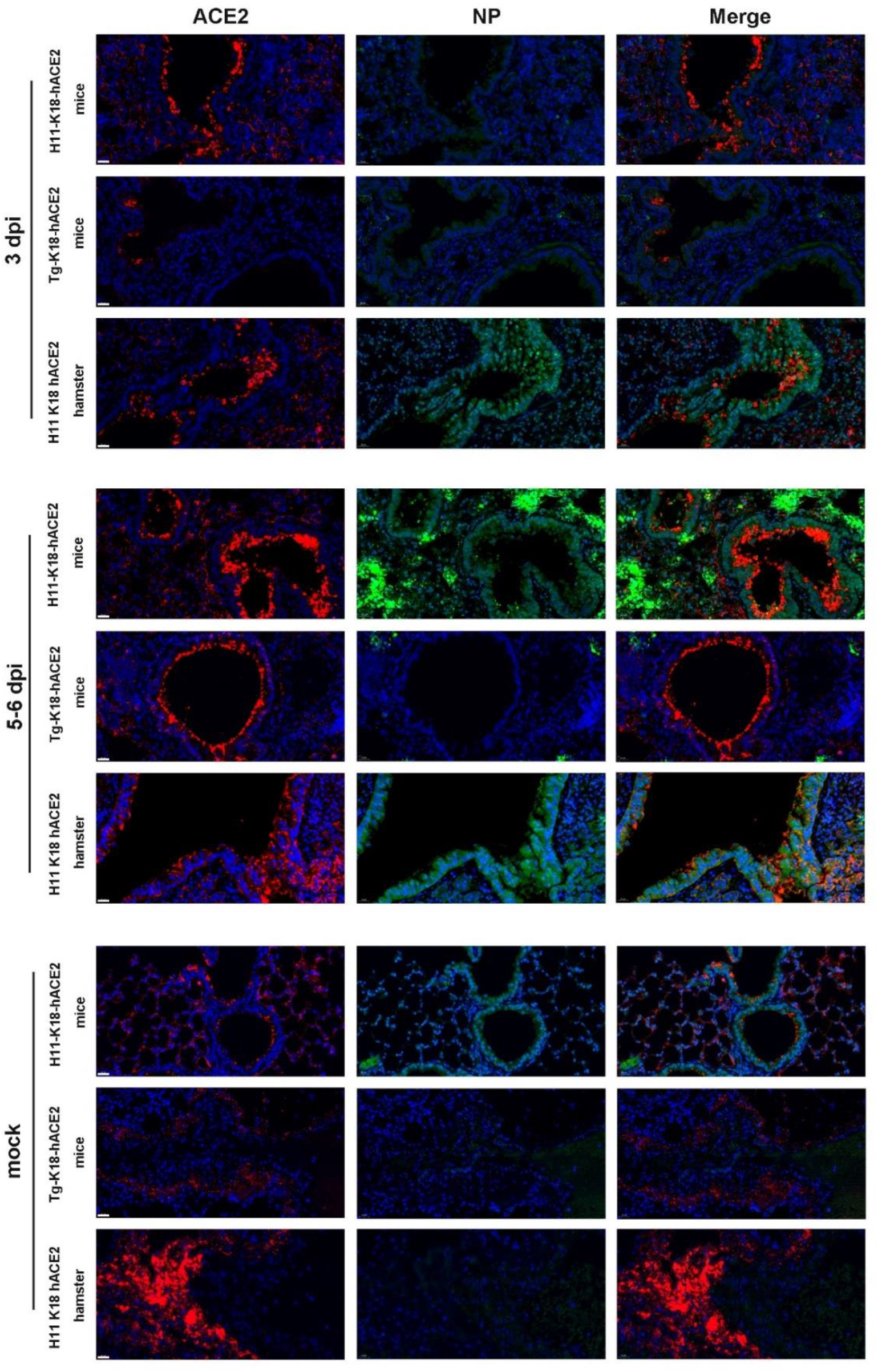
Immunofluorescence analysis in the lungs of mice and hamsters inoculated with SARS-CoV-2 BA.5 variant. Euthanized mice and hamsters (3-4 each) were used to examine the SARS-CoV-2 NP protein and ACE2 receptor expression in the lungs at 3 dpi. Natural death mice and hamsters (6 each) were used to examine the SARS-CoV-2 NP protein and ACE2 receptor expression in the lungs 5-6 dpi.

**Fig. 6.**
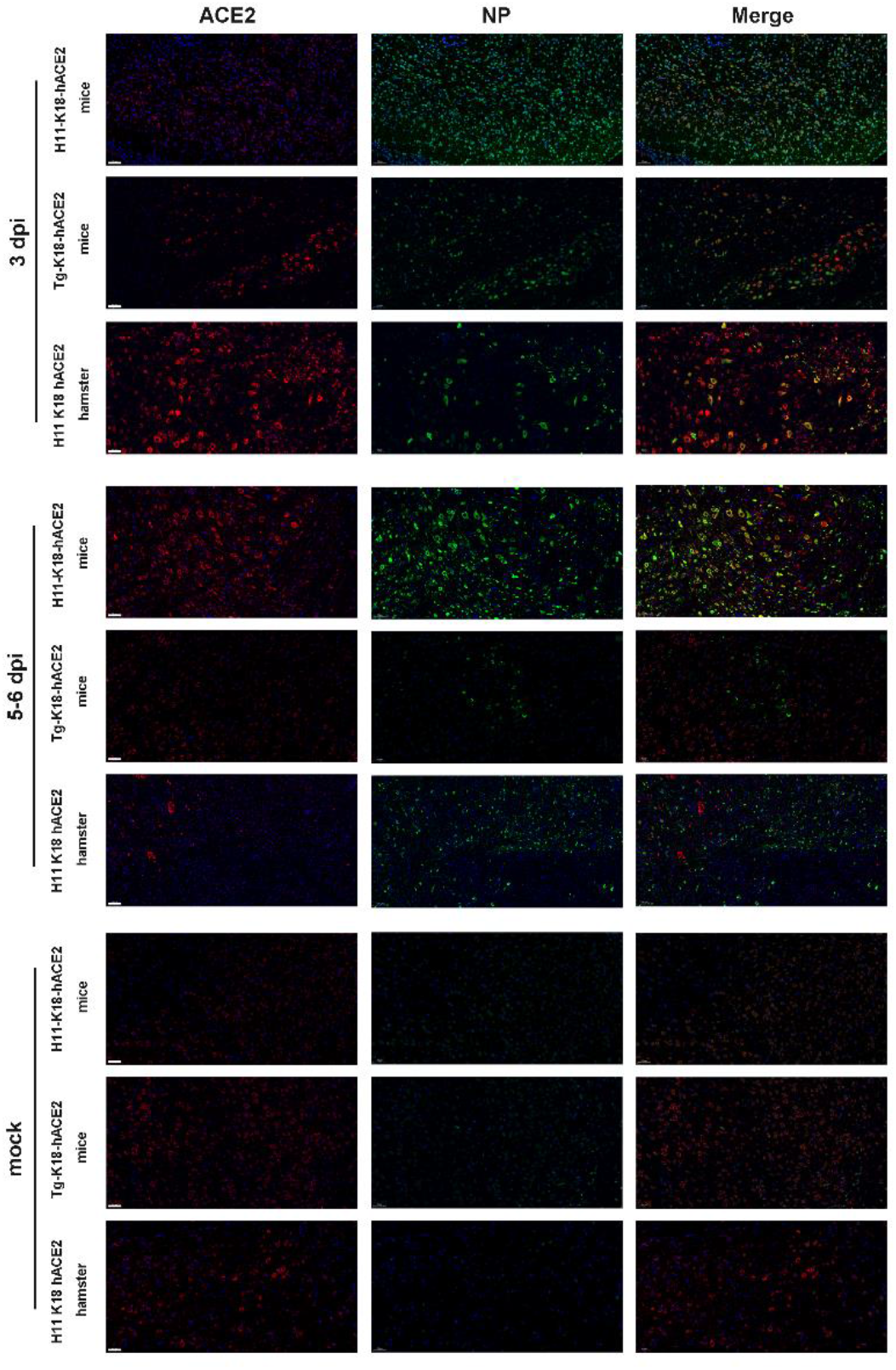
Immunofluorescence analysis in the brains of mice and hamsters inoculated with SARS-CoV-2 BA.5 variant. Euthanized mice and hamsters (3-4 each) were used to examine the SARS-CoV-2 NP protein and ACE2 receptor expression in the brains 3 dpi. Natural death mice and hamsters (6 each) were used to examine the SARS-CoV-2 NP protein and ACE2 receptor expression in the brains 5-6 dpi.

Furthermore, a concurrent upregulation of ACE2 expression was observed in the brains of all mice and hamsters during SARS-CoV-2 invasion, as compared to mock models[22].

## Discussion

Our investigation has yielded significant insights into establishing an animal model with heightened susceptibility to SARS-CoV-2 infection through the introduction of ectopic hACE2 expression in Syrian hamsters. This model manifests a severe and ultimately fatal disease phenotype, thereby corroborating and extending the findings of preceding studies focused on SARS-CoV-2 infection dynamics[18,23,24]. Notably, our experimentation with BA.5 variants has unveiled a distinctive viral dissemination pattern, encompassing critical organs such as the lungs and brain, ultimately resulting in mortality.

Of particular interest is the comparison between the hamster model, specifically the K18-hACE2 hamsters, and the corresponding mouse models, including H11-K18-hACE2 mice and Tg-K18-hACE2 mice. Remarkably, K18-hACE2 hamsters exhibited exacerbated lung injury compared to their murine counterparts. This disparity underscores the importance of employing a suitable animal model for accurate disease characterization. Moreover, despite previous reports suggesting alternative outcomes, our study has brought to light the lethality associated with BA.5 infection in all three models – H11-K18-hACE2 mice, Tg-K18-hACE2 mice, and H11-K18-hACE2 hamsters. This lethality mirrors the severe clinical manifestations of COVID-19 observed in human patients, reinforcing the pertinence of our chosen models in the study of the distinct SARS-CoV-2 BA.5 variants. While the brains of animal models exhibited no discernible pathological variances, it is notable that the expression of NP in mouse and hamster models indicated that hamster models offer a closer approximation of human physiology[25].

Hamster models have gained widespread acceptance within the scientific community as valuable tools for assessing therapeutic interventions against SARS-CoV-2 and comprehensively characterizing the disease[9,23,24,26,27]. To ensure robust comparisons, our study simultaneously subjected three distinct models to BA.5 infection. This approach facilitated the elucidation of nuanced distinctions between these models, ultimately highlighting the hamster model’s superior fidelity in capturing the characteristics of the SARS-CoV-2 BA.5 variants. The hamsters showed similar patterns in histopathological analysis compared to humans late in the illness[28]. Even though hamsters showed more lethality when infected with the disease, the reason probably has to do with the differences in the size of the brain and lungs.

Nonetheless, it is essential to acknowledge the inherent limitations of our study. Foremost, the absence of immunological validation raises the possibility of gaps in the comprehensiveness of our conclusions. Furthermore, the precise mechanism underpinning the lethality induced by the BA.5 variants remains elusive, a challenge not uncommon in infection-based research. Given these limitations, we advocate for continued investigations aimed at addressing these gaps and unraveling the intricate mechanisms that drive the observed outcomes. This pursuit promises to enhance our understanding of SARS-CoV-2 infection dynamics and potentially inform novel therapeutic strategies. For future studies, we could prioritize the categorization of hamster models, particularly examining how aging may influence lethality among H11-K18-hACE2 hamsters[11,29,30]. The detailed investigation is fundamental for enhancing our understanding of COVID-19, tailoring treatments, and optimizing experimental methodologies for more effective research outcomes.

## Materials and Methods

### Viruses and cells

The SARS-CoV-2 BA.5 variant was acquired from the National Virus Resource Center at the Wuhan Institute of Virology. All virus-related work was conducted within a Biosafety Level 3 (BSL-3) containment facility at the Laboratory Animal Center of the Wuhan Institute of Virology, CAS. Regular testing of cells consistently yielded negative results for mycoplasma using a PCR-based assay. The Cercopithecus aethiops kidney cells (Vero E6, ATCC® CRL-1586) were used for virus propagation and titration. Vero E6 cells were cultured in DMEM supplemented with 10% fetal bovine serum (FBS, Gibco) and 1% Anti-Anti (Invitrogen, USA) at 37°C in a 5% CO2 environment.

### Animal models

Golden hamsters were procured from the Charles River Company in Peking, China, and were maintained under a 14-hour light/10-hour dark cycle at the Animal Facility Center of Nanjing Medical University. The H11-pK18-hACE2 transgenic mice were obtained from Gempharmatech Co., Ltd (Nanjing, China). The Nanjing Medical University Institutional Animal Care and Research Committee granted ethical approval for gene editing experiments (IACUC code, 2108028). Moreover, all experiments involving virus infection in animals were approved by the Institutional Animal Care and Use Committee (approval number: WIVA05202204) at the Wuhan Institute of Virology. All animal experiments concerning virus infection were conducted in an animal biosafety level 3 laboratory at the Wuhan Institute of Virology.

### Plasmid

In terms of plasmids, Supplementary Table 1 provides the sequences of all oligonucleotides used. The oligonucleotides were synthesized at GenScript Biotech Corporation. The pK18-hACE2 plasmid was obtained from Addgene (Cat No:149449) (31). To create the gH11-PB3-pK18-hACE2-PB5 plasmid, the first step involved generating a gH11 homologous pUC57-H11-mcs plasmid with a multiple cloning site. Approximately 1000 bp of the homologous arm of the H11 site was amplified by PCR from golden hamster genomic DNA and inserted into the multiple cloning site of the pUC57 vector. In the second step, the PB 3’ and 5’ inverted terminal repeats (ITRs) were introduced into the pUC57-H11-mcs plasmid to create the PB transposon vector, pUC57-gH11-PB3-mcs-PB5. Finally, pK18-hACE2 fragments were cut using HpaI and SalI enzymes and inserted into the multiple cloning site of the pUC57-gH11-PB3-mcs-PB5 plasmid.

### Generation of H11-K18-hACE2 trangenic golden hamsters

The process of generating H11-K18-hACE2 transgenic golden hamsters involved designing H11-SgRNAs for golden hamsters using CRISPRdirect (http://crispr.dbcls.jp/) and CRISPOR (http://crispor.tefor.net/). The complementary target oligonucleotides were cloned into the pUC57-T7-sgRNA-trcRNA vector through the BsaI restriction site. The sequences containing T7 promoter and sgRNA were PCR-amplified and used as templates to produce sgRNAs through in vitro transcription using the HiScribe T7 High Yield RNA Synthesis Kit (NEB, E2040S). All sgRNAs were purified by lithium chloride precipitation and stored at −80°C.

For generating genome-modified golden hamsters, as described (32), CRISPR– Cas9 protein (IDT), sgRNA, and gH11-PB3-pK18-hACE2-PB5 vector were injected into two-cell embryos in a red-light room under a microscope with a red filter. The injected embryos were cultured in mHECM9+HSA medium in a humidified 10% CO2 environment at 37.5°C for 15 –30 min and then transferred to the oviducts of 0.5-day pregnant recipients, with 15–30 embryos for each recipient. Founder pups were mated with WT males and females to produce the F1 generation. Genotyping of F1 pups was performed using the one-step mouse genotyping kit according to the manufacturer’s instructions (Vazyme, PD101–01). A list of all oligo sequences is provided in Supplementary Table 1.

### Generation of K18-hACE2 transgenic mice

Circular gH11-PB3-pK18-hACE2-PB5 plasmids at a concentration of 3 ng/μl were mixed with Super PiggyBac Transposase plasmid at a concentration of 1.5 ng/μl (Supplementary Figure 1). The DNA samples (5 ng/μl) were microinjected into fertilized C57B6 oocytes by IVF. Genotyping of pups was performed as described above.

### Mice and golden hamster infection

The animals were anesthetized with tribromoethanol (Avertin) (250 mg/kg) and intranasally infected with 1∗104 TCID50 (for naive infection) SARS-CoV-2 BA.5 virus in 50 μL DMEM per animal. The experiments were divided into three groups: the control group (n=3) inoculated with DMEM, and the pathology progression group inoculated with the virus (n=10) (Table S2). Accidental deaths or uninfected mice were excluded from subsequent experiments. Mice were weighed and observed for clinical signs daily throughout the study. Four animals were euthanized at 3, 5, 6, or 7 DPI, and mice presenting more than a twenty percent decrease in body weight were humanely euthanized and considered deceased animals. The details regarding the number of animals in each group are shown in Table S3. Tissues, including blood, heart, liver, spleen, lung, brain, kidney, testis, and small intestines, were harvested.

### Determination of virus titers in tissues

Vero-E6 cells were seeded in 24-well plates 1 day before use. Infected tissues were homogenized in DMEM. The cells were inoculated with tissue dilutions for 1 h. Next, the inoculum was removed, and the cells were incubated with 0.9% methylcellulose for 5 days. Plaque-forming units (PFU) were counted after crystal violet staining to calculate the viral titer.

### Histological analysis

Samples were fixed with 4% paraformaldehyde, paraffin-embedded, and cut into 3.5-μm sections. Fixed tissue samples were used for hematoxylin-eosin (H&E) staining and indirect immunofluorescence assays (IFA) for the detection of the SARS-CoV-2 antigen. Routine histology involved staining tissue sections with H&E.

For IFA, slides were deparaffinized and rehydrated, followed by 15-min heat-induced antigen retrieval with EDTA pH 8.0 in a microwave oven. The slides were washed with PBS/0.02% Triton X-100, then blocked with 5% BSA at room temperature for 1 h. A primary antibody (rabbit anti-SARS-CoV-2 N protein polyclonal antibody, 1:500, generated in Shi ZL’s lab; and mouse ACE2 antibody, Servicebio, 1:1000) was added dropwise to the sections and then washed with PBS. After the slides were slightly dried, tissues were covered with Cy3-conjugated goat-anti-rabbit IgG (Abcam, ab6939) at a 1:200 dilution. Following PBS wash, slides were stained with DAPI (Beyotime) at a 1:100 dilution. For Martius Scarlet Blue (MSB) staining, fixed tissue sections were stained with hyposulfite solution, Celestine blue, and hematoxylin for 3-5 minutes, respectively. After acidic differentiation with acid alcohol, the sections were stained with Martius Yellow solution, carmoisine solution, phosphotungstic acid solution, and aniline blue staining solution, respectively. Tissue sections were rinsed with 1% acetic acid and then dehydrated, cleared, and mounted with neutral gum. Image information was collected using a Pannoramic MIDI system (3DHISTECH, Budapest) and FV1200 confocal microscopy (Olympus).

### Western blot analysis

For Western blot analysis, tissues for western blot were obtained from golden hamsters (aged 8-12 weeks). Total proteins were extracted in 95% Laemmle sample buffer, separated on 10% SDS–PAGE using the Mini-PROTEAN Tetra Cell System (Bio-Rad), and electrophoretically transferred to polyvinylidene difluoride membranes. Anti-β-actin (ACTB) mouse monoclonal antibodies (Proteintech, 66009-1-lg, 1:2,000) and anti-ACE2 rabbit polyclonal antibodies (1:1,000) were used as primary antibodies and incubated with the membranes at 4°C for 12 hours. After washing twice with PBS containing 0.2% Tween-20 (PBST), the membranes were incubated with HRP-linked goat anti-rabbit antibodies (ThermoFisher Scientific, 31466, 1:5,000) or HRP-linked anti-mouse IgG (ThermoFisher Scientific, 31430, 1:5,000) at room temperature for 1 hour before visualization using the ECL Detection kit (GE Healthcare).

### Statistical Analysis

Statistical analyses were performed using a two-tailed Student’s t-test or two-way ANOVA in GraphPad Prism 8. Data were expressed as means ± SEM. Statistical significant: *p < 0.05, ** p < 0.01, *** p < 0.001, **** p < 0.0001

## Acknowledgments

We thank all members of J.M Li’s and Shi ZL laboratories for their discussion and comments on this project; We thank all members from the animal center of the Nanjing Medical University for the transgenic mice and golden hamster breeding; We thank Tao Du and Jin Xiong from the Center for Biosafety Mega-Science of the Wuhan Institute of Virology for their essential support.

J.M L, A.M S, ZL S conceived and designed the study. W.T Z. generated the H11-K18-hACE2 knockin golden hamster and Tg(K18-hACE2) transgenic mice with the help of Y.N L, D.M and A.H Z, D.M,M.H W and Y. G analyzed the phenotype of the H11-K18-hACE2 knockin golden hamster and Tg(K18-hACE2) transgenic mice with help of Y.N L.H.F L, M.Q L, and R.D J performed virus culturing experiments.. J.M L, D. M, H.F L, B. H, and M. P interpreted the data of the experiments and wrote the manuscript. Writing – Original Draft, Y.G, M.D, M.P Writing – Review & Editing, Shi Z L and J.M L.

## Data Availability

The datasets generated during and/or analyzed during the current study are available from the corresponding author on reasonable request.

---

**Supplementary Figure 1:**
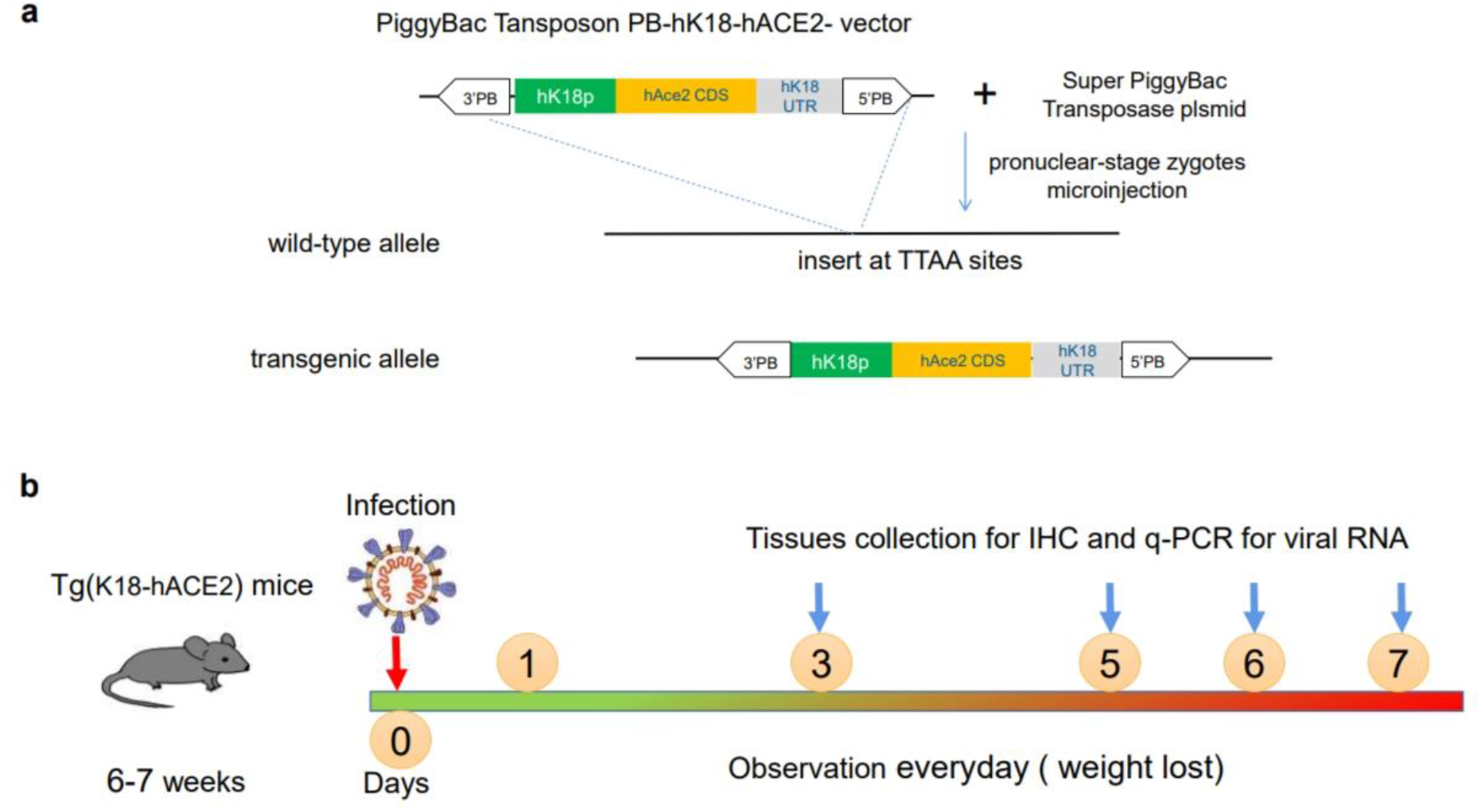

**Supplementary Table 1:**
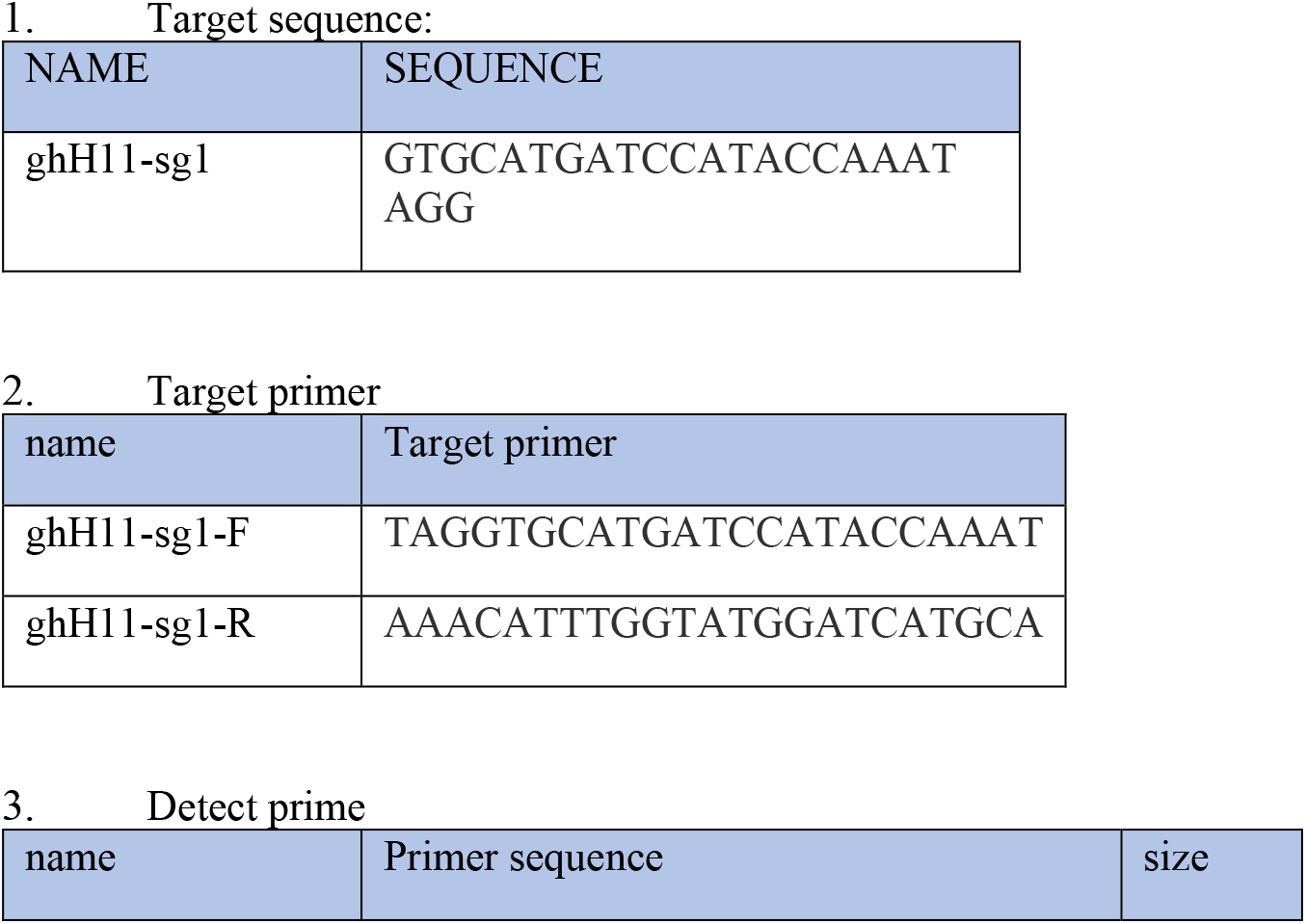
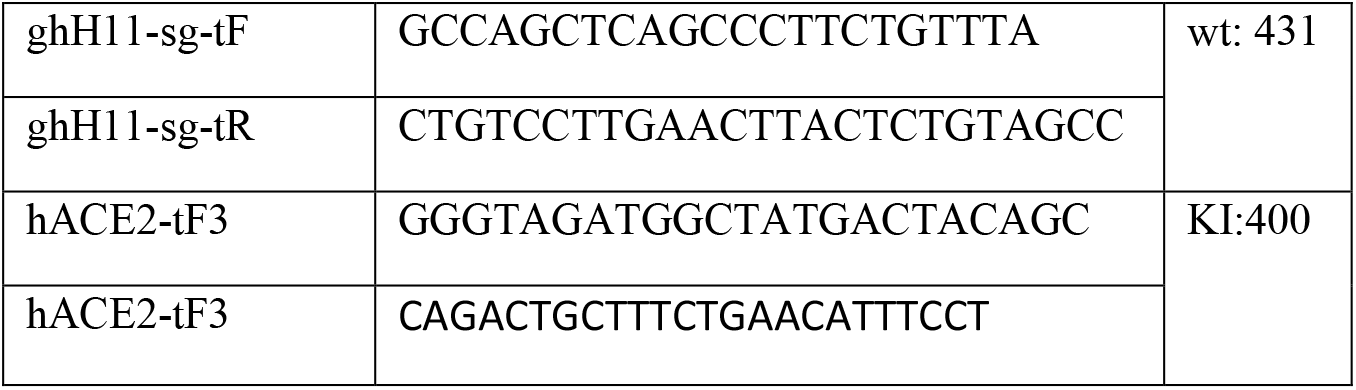
Table 1 all primers for this paper

## References

1. Chen N, Zhou M, Dong X, Qu J, Gong F, Han Y, et al. Epidemiological and clinical characteristics of 99 cases of 2019 novel coronavirus pneumonia in Wuhan, China: a descriptive study. The Lancet. 2020;395:507–13.

2. Zhou P, Yang X-L, Wang X-G, Hu B, Zhang L, Zhang W, et al. A pneumonia outbreak associated with a new coronavirus of probable bat origin. Nature. 2020;579:270–3.

3. Hachmann NP, Miller J, Collier AY, Ventura JD, Yu J, Rowe M, et al. Neutralization Escape by SARS-CoV-2 Omicron Subvariants BA.2.12.1, BA.4, and BA.5. New England Journal of Medicine. 2022;387:86–8.

4. Tuekprakhon A, Nutalai R, Dijokaite-Guraliuc A, Zhou D, Ginn HM, Selvaraj M, et al. Antibody escape of SARS-CoV-2 Omicron BA.4 and BA.5 from vaccine and BA.1 serum. Cell. 2022;185:2422-2433.e13.

5. Cao Y, Yisimayi A, Jian F, Song W, Xiao T, Wang L, et al. BA.2.12.1, BA.4 and BA.5 escape antibodies elicited by Omicron infection. Nature. 2022;608:593–602.

6. Wan Y, Shang J, Graham R, Baric RS, Li F. Receptor Recognition by the Novel Coronavirus from Wuhan: an Analysis Based on Decade-Long Structural Studies of SARS Coronavirus. J Virol. 2020;94.

7. Gretebeck LM, Subbarao K. Animal models for SARS and MERS coronaviruses. Curr Opin Virol. 2015;13:123–9.

8. Golden JW, Cline CR, Zeng X, Garrison AR, Carey BD, Mucker EM, et al. Human angiotensin-converting enzyme 2 transgenic mice infected with SARS-CoV-2 develop severe and fatal respiratory disease. JCI Insight. 2020;5.

9. Chan JF-W, Zhang AJ, Yuan S, Poon VK-M, Chan CC-S, Lee AC-Y, et al. Simulation of the Clinical and Pathological Manifestations of Coronavirus Disease 2019 (COVID-19) in a Golden Syrian Hamster Model: Implications for Disease Pathogenesis and Transmissibility. Clinical Infectious Diseases. 2020;

10. Halfmann PJ, Iida S, Iwatsuki-Horimoto K, Maemura T, Kiso M, Scheaffer SM, et al. SARS-CoV-2 Omicron virus causes attenuated disease in mice and hamsters. Nature. 2022;603:687–92.

11. DeGrace MM, Ghedin E, Frieman MB, Krammer F, Grifoni A, Alisoltani A, et al. Defining the risk of SARS-CoV-2 variants on immune protection. Nature. 2022;605:640–52.

12. Chu H, Chan JF-W, Yuen K-Y. Animal models in SARS-CoV-2 research. Nat Methods. 2022;19:392–4.

13. Gilliland T, Dunn M, Liu Y, Alcorn MDH, Terada Y, Vasilatos S, et al. Transchromosomic bovine-derived anti-SARS-CoV-2 polyclonal human antibodies protects hACE2 transgenic hamsters against multiple variants. iScience. 2023;26:107764.

14. Hippenmeyer S, Youn YH, Moon HM, Miyamichi K, Zong H, Wynshaw-Boris A, et al. Genetic Mosaic Dissection of Lis1 and Ndel1 in Neuronal Migration. Neuron. 2010;68:695–709.

15. Tasic B, Hippenmeyer S, Wang C, Gamboa M, Zong H, Chen-Tsai Y, et al. Site-specific integrase-mediated transgenesis in mice via pronuclear injection. Proceedings of the National Academy of Sciences. 2011;108:7902–7.

16. Zheng J, Wong L-YR, Li K, Verma AK, Ortiz ME, Wohlford-Lenane C, et al. COVID-19 treatments and pathogenesis including anosmia in K18-hACE2 mice. Nature. 2021;589:603–7.

17. Oladunni FS, Park J-G, Pino PA, Gonzalez O, Akhter A, Allué-Guardia A, et al. Lethality of SARS-CoV-2 infection in K18 human angiotensin-converting enzyme 2 transgenic mice. Nat Commun. 2020;11:6122.

18. Uraki R, Halfmann PJ, Iida S, Yamayoshi S, Furusawa Y, Kiso M, et al. Characterization of SARS-CoV-2 Omicron BA.4 and BA.5 isolates in rodents. Nature. 2022;612:540–5.

19. Tamura T, Yamasoba D, Oda Y, Ito J, Kamasaki T, Nao N, et al. Comparative pathogenicity of SARS-CoV-2 Omicron subvariants including BA.1, BA.2, and BA.5. Commun Biol. 2023;6:772.

20. Zhang L, Zhou L, Bao L, Liu J, Zhu H, Lv Q, et al. SARS-CoV-2 crosses the blood–brain barrier accompanied with basement membrane disruption without tight junctions alteration. Signal Transduct Target Ther. 2021;6:337.

21. Puelles VG, Lütgehetmann M, Lindenmeyer MT, Sperhake JP, Wong MN, Allweiss L, et al. Multiorgan and Renal Tropism of SARS-CoV-2. New England Journal of Medicine. 2020;383:590–2.

22. Castaneda DC, Jangra S, Yurieva M, Martinek J, Callender M, Coxe M, et al. Spatiotemporally organized immunomodulatory response to SARS-CoV-2 virus in primary human broncho-alveolar epithelia. iScience. 2023;26:107374.

23. Davies M-A, Morden E, Rousseau P, Arendse J, Bam J-L, Boloko L, et al. Outcomes of laboratory-confirmed SARS-CoV-2 infection during resurgence driven by Omicron lineages BA.4 and BA.5 compared with previous waves in the Western Cape Province, South Africa. International Journal of Infectious Diseases. 2023;127:63–8.

24. Wolter N, Jassat W, Walaza S, Welch R, Moultrie H, Groome MJ, et al. Clinical severity of SARS-CoV-2 Omicron BA.4 and BA.5 lineages compared to BA.1 and Delta in South Africa. Nat Commun. 2022;13:5860.

25. Imbiakha B, Ezzatpour S, Buchholz DW, Sahler J, Ye C, Olarte-Castillo XA, et al. Age-dependent acquisition of pathogenicity by SARS-CoV-2 Omicron BA.5. Sci Adv. 2023;9.

26. Bryche B, St Albin A, Murri S, Lacôte S, Pulido C, Ar Gouilh M, et al. Massive transient damage of the olfactory epithelium associated with infection of sustentacular cells by SARS-CoV-2 in golden Syrian hamsters. Brain Behav Immun. 2020;89:579–86.

27. de Melo GD, Lazarini F, Levallois S, Hautefort C, Michel V, Larrous F, et al. COVID-19– related anosmia is associated with viral persistence and inflammation in human olfactory epithelium and brain infection in hamsters. Sci Transl Med. 2021;13.

28. Roberts A, Vogel L, Guarner J, Hayes N, Murphy B, Zaki S, et al. Severe Acute Respiratory Syndrome Coronavirus Infection of Golden Syrian Hamsters. J Virol. 2005;79:503–11.

29. McCray PB, Pewe L, Wohlford-Lenane C, Hickey M, Manzel L, Shi L, et al. Lethal Infection of K18-hACE2 Mice Infected with Severe Acute Respiratory Syndrome Coronavirus. J Virol. 2007;81:813–21.

30. Zhang H, Zhang F, Chen Q, Li M, Lv X, Xiao Y, et al. The piRNA pathway is essential for generating functional oocytes in golden hamsters. Nat Cell Biol. 2021;23:1013–22.

