## Supplemental Table2 for "Characterization of the SARS-CoV-2 BA.5 Variants in H11-K18-hACE2 Hamsters"

**Table S1.**

1. Target sequence:

| NAME | SEQUENCE |
|---|---|
| ghH11-sg1 | GTGCATGATCCATACCAAAT AGG |

2. Target primer

| name | Target primer |
|---|---|
| ghH11-sg1-F | TAGGTGCATGATCCATACCAAAT |
| ghH11-sg1-R | AAACATTTGGTATGGATCATGCA |

3. Detect prime

| name | Primer sequence | size |
|---|---|---|
| ghH11-sg-tF | GCCAGCTCAGCCCTTCTGTTTA | wt: 431 |
| ghH11-sg-tR | CTGTCCTTGAACTTACTCTGTAGCC | |
| hACE2-tF3 | GGGTAGATGGCTATGACTACAGC | KI:400 |
| hACE2-tF3 | CAGACTGCTTTCTGAACATTTCCT | |

**Primers used in this paper.**
